# Generative World Models to compute protein folding pathways

**DOI:** 10.1101/2025.03.26.645554

**Authors:** Alan Ianeselli, Jewon Im, Eddie Cavallin, Mark B. Gerstein

## Abstract

We developed a generative World Model in the context of biomolecular simulations for proteins. An agent was trained by evolution strategy in a latent spatio-temporal representation of the structural transitions, to learn a policy to drive protein folding simulations via dihedral rotations (Φ, Ψ). Latent configurations were decoded back into atomistic structures using atomistic force fields, allowing for the calculation of structural properties and energetic terms. We show how the model can be regularized by adding Ramachandran-based rewards during the training of the controller. Results have been validated against equilibrium MD data. Markov State Modeling could be applied to reconstruct the dynamics of the system and reconstruct the unbiased folding landscape. The method proposed here facilitates the study of the conformational changes and folding pathways of proteins by generative AI, and can have application in structure-based drug discovery.

## INTRODUCTION

AI models often “hallucinate,” meaning they generate information that is not grounded in real data but instead synthesized from learned patterns, inventing new scenarios on their own^1^. While this is often a drawback for many tasks that require strict factual accuracy, the phenomenon can be exploited in the so-called “World Models”^2^. They are a class of generative neural networks designed to simulate environments for training reinforcement learning agents. World Models create a latent spatial and temporal representation of an environment - sometimes referred to as a “dream environment” - in which an AI agent can explore, learn, and optimize decision-making policies in the latent space instead of the real environment. By leveraging this AI’s ability to hallucinate possible future states, World Models allow reinforcement learning agents to develop and refine strategies in a controlled, imagined space before being deployed in the real environment, strongly reducing the computational complexity during training. The original World Model implementation from Ha and Schmidhuber^2^ was developed to teach an agent model to solve a car racing task, or learn to avoid fireballs shot by monsters. The model was compressing visual information into a latent spatiotemporal representation. Later models were developed for other tasks such as games like Atari^3^, Go^4^, Chess^4^, as well as autonomous driving^5^, video generation^6^ and robotics^7^.

In this paper we develop, for the first time, a generative World Model to evolve a policy to compute protein folding pathways. Protein folding is a high-dimensional problem characterized by a vast conformational space, where a protein can adopt an astronomical number of possible configurations^8^. The process is governed by intricate energetic landscapes, complex atomistic interactions, and stochastic fluctuations, making it a highly nonlinear and computationally intensive process to study.

Efficient optimization methods such as enhanced path sampling techniques and AI-based methods^9–12^, are often necessary over equilibrium approaches, such as Molecular Dynamics^13^, in order to explore folding mechanisms within feasible timeframes and computational resources. Atomistic protein simulations are very computationally intensive, due to the high number of atoms and degrees of freedom of the system. Recently, with the advent of generative AI, some studies tackled this challenge by making use of Variational Autoencoders (VAEs) to reduce the dimensionality, and project the high-dimensional protein structure onto low-dimensional latent vectors. This facilitated the study of protein structural ensembles^14,15^, conformational changes^16,17^, and disordered proteins^18,19^.

The method we propose here is based on Monte Carlo protein simulations^20^, where random torsions of the backbone dihedrals (Φ and Ψ) take place to explore the conformational landscape of proteins. First, the dimensionality of the protein configurations is reduced by a Variational Autoencoder into smaller latent vectors. Then, a small Feed Forward Neural Network (FFN) is trained to predict the next latent protein configuration given the current latent configuration and a latent dihedral rotation (action) performed on it. In this way, the FFN has built its own understanding of the structural transitions in the folding “dream” environment. Finally, a small controller is trained in this dream environment to perform the necessary dihedral rotations in order to drive the folding towards the target state (i.e. the native state). The decoder of the VAE can then be used to reconstruct the full atomistic representation of the protein configurations.

It would be challenging to train our agent in the atomistic environment, because the high dimensionality of the parameter space would make the training computationally very demanding. Instead, the training was performed in the dream environment - a compressed (latent) spatiotemporal representation of the structural transitions. The controller learned to perform the latent rotations on the protein dihedral angles Φ and Ψ, driving the conformational transitions towards the target of interest (i.e. the folded state).

The method has been applied to compute the folding pathway of TrpCage miniprotein (PDB: 1L2Y, 20 aa), Fip35 WW domain (PDB: 2F21, 35 aa), Protein G (PDB: 1MI0, 56 aa) and human Cellular Retinol Binding Protein II (PDB: 4ZGU, 133 aa), and to calculate their folding landscapes. We show how the model can be regularized using a Ramachandran-based score function during the policy evolution, discouraging structures with unrealistic Φ and Ψ combinations. This yielded folding trajectories that followed lower energy pathways. Our method was also compared to other enhanced path sampling methods and, in some cases, showed good accuracy. The method, however, yields folding trajectories and structures that are out-of-equilibrium and sometimes unrealistic. Performing short relaxations of the structures through MD allowed for the reconstruction of the unbiased dynamics of the proteins via Markov State Modeling (MSM).

This method is a strategy to compute the folding pathway of biomolecules and obtain the atomistic structure of the intermediate states using state-of-the-art atomistic force fields. It can have applications in Structure-Based Drug Discovery (SBDD), to develop pharmacological targeting strategies against structures along the folding pathways^21^, as well as to elucidate the biological role of folding intermediates and the atomistic details of the conformational transitions.

### Model and training procedure

Building this generative World Model involved the following steps:

-<u>Step 1</u>: collect random rollout data. Starting from arbitrary protein conformations (the native or randomly generated unfolded states), we collected trajectories with random dihedral moves (i.e. random Φ and Ψ torsions ranging between 0 and ± 30°). After each move, an energy minimization step was performed to relax the structure and eliminate atomic clashes. Structures have been generated with pyRosetta (Python-based interface to the Rosetta molecular modeling tool^22^), using a state-of-the-art atomistic force field (ref2015^23^). For each protein, the training set corresponded to 50 trajectories of 500 steps each, for a total of 25000 protein configuration and actions. The validation and test set consisted of 5 additional trajectories of 500 steps each (total 2500 protein configurations and actions each). This training set contains the information on how different dihedral rotations change the structure of the protein.

-<u>Step 2</u>: train Variational AutoEncoder 1 (VAE1). The rollout data were used to train VAE1 to encode the protein features (a concatenation of Φ dihedrals, Ψ dihedrals and C_α_ distance from the native state) into a lower dimensional vector that we named *z*_features_ of length 10. VAE1 is therefore responsible for lowering the dimensionality of the structures into lower dimensional vectors, strongly reducing the number of parameters. The decoder of VAE1 can later be used to reconstruct back the full atom structure from the latent vector z_features_.

-<u>Step 3</u>: train Variational AutoEncoder 2 (VAE2). The rollout data were used to train VAE2 to encode the action (ΔΦ and ΔΨ) into a latent vector that we named *z*_action_, also of length 10. VAE2 is therefore responsible for reducing the number of dimensions required to describe the action. The VAEs of steps 2 and 3 ensure that the features are mapped in the latent space continuously. This is important in our context of protein folding, because we want to be able to draw realistic and configurations when decoding from the latent space back into the real environment. The loss function for both VAE1 and VAE2 consisted in the reconstruction loss (MAE, Mean Average Error) between the features (after decoding) and the Kullback-Leibner divergence^24^ term to ensure that the latent space follows a standard normal distribution. Additional information in supplementary section 4.

-<u>Step 4</u>: train the Feed Forward Model (FFN) model to predict the next latent configuration given the current latent configuration and the latent action performed on it (*z*_features_(*t*+1) | *z*_features_(*t*), *z*_action_(*t*)). The FFN is the core of this World Model. It learns how an action affects the current configuration and predicts the next one, entirely in the latent space. First of all, the trained encoders of VAE1 and VAE2 are used to generate the training data for the FFN, i.e. calculating the z_features_ and z_action_ of the rollout. Then, the data are used to train FFN. The loss function of the FFN consisted of the reconstruction error (MAE) between the true and the predicted *z*_features_(*t*+1).

-<u>Step 5</u>: train the controller to predict the action to be performed on the given current configuration (*z*_action_ | *z*_features_), in order to drive the trajectory closer to the target (i.e. the folded state), starting from an arbitrary unfolded state. The trained FFN is used to predict the next z_features_(*t*+1) given the current z_features_(*t*) and the z_action_(*t*) output of the controller. The controller was trained using the evolutionary strategy CMA (Covariance Matrix Adaptation^25^). CMA is a powerful evolutionary optimization algorithm that is particularly well-suited for training controllers in complex, high-dimensional search spaces. For protein folding, credit assignment - the process of determining which parameters contributed most to a successful outcome - is highly nontrivial. CMA explores solution spaces through evolutionary strategies, refining a population of candidate solutions over successive generations. Note how the training of the controller is performed entirely in the latent space and in the FFN’s own understanding of the environment (aka “in-dream” training). At each generation, each agent (population size of 20) performs 80 or 100 iterations starting from the unfolded z_features_. The reward to the agent corresponded to the distance (mean average error, MAE) between the last latent configuration (*z*_features_(*t*+100)) and the native latent configuration, plus a Ramachandran-based regularization term (further details in the later section). For simplicity, the objective of the training was to minimize the reward. The overall architecture is shown in Figure 1a-b. Additional model details can be found in the supplementary information section 1.

**Figure 1.**
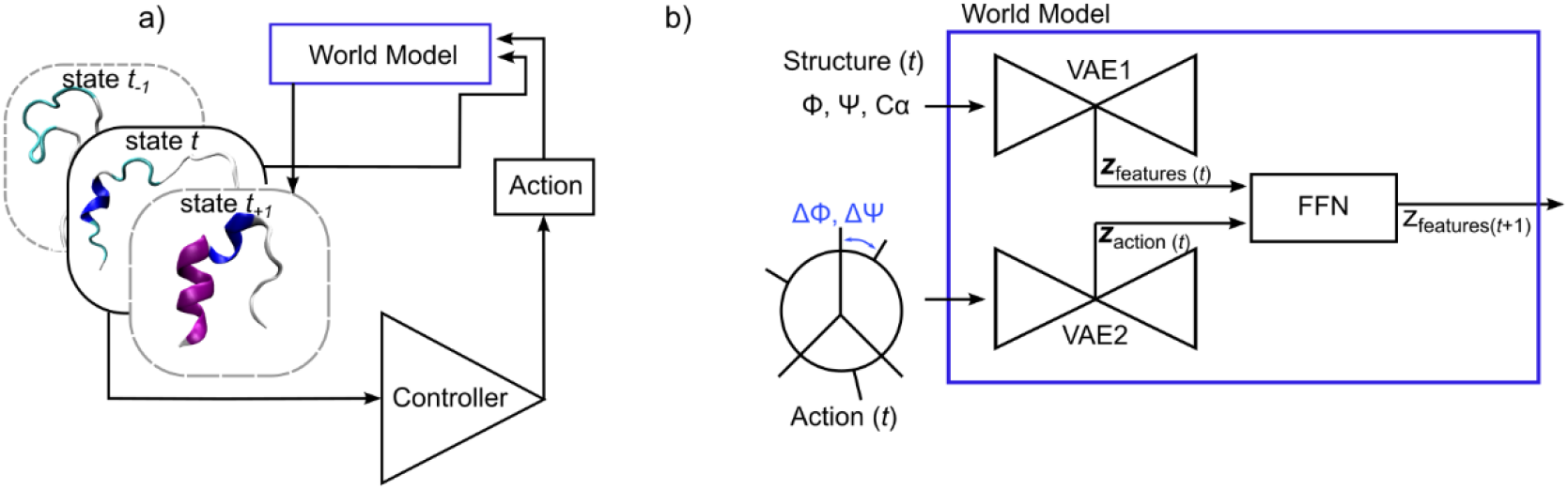
Generative World Model to compute protein folding pathways. a) The World Model is employed in a reinforcement learning framework, where a controller is trained to predict actions to be performed on proteins to drive their folding and generate the next structure. b) Architecture of the World Model. It is composed of two Variational AutoEncoders (VAE1, VAE2) that encode the protein structure (Φ, Ψ dihedrals and C_α_ distances from native) and the action (ΔΦ and ΔΨ) into latent vectors. A Feed Forward Network (FFN) predicts the next latent structure given the current latent structure and action.

The trained model is then employed for the inference of folding trajectories, starting from an arbitrary unfolded conformation and predicting the conformational changes that lead to the target state (in our case, the folded state). The atomistic structures at each step can be reconstructed using the decoder of VAE1. The reconstruction procedure is indicated in the supplementary section 3.

## RESULTS

### Regularization

To reduce overfitting during policy evolution, we have introduced a path-dependent Ramachandran-based energy score as reward during the training of the controller. The Ramachandran score is calculated on the normalized frequency distribution of the protein backbone dihedral angles Φ, Ψ, computed from ∼8000 high resolution protein PDBs (obtained from^26,27^). The Ramachandran score discourages structures with unrealistic Φ and Ψ combinations, which would be sterically unfavorable.

During controller training, the z_features_ of each step was decoded back using the decoder from VAE1 in order to reconstruct the values of Φ and Ψ and calculate the Ramachandran-based energy score. The total reward, in our case to be minimized, can be defined as following:

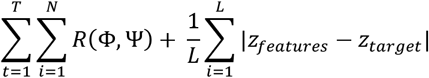

where *R*(Φ,Ψ) corresponds to the Ramachandran matrix shown in Figure 2a, summed over *N* protein residues for every frame *t* of the trajectory of length *T*. The second term corresponds to the MAE between the target latent *z* vector (*z*_*target*_) and the last *z* vector of the trajectory (*z*_*features*_), with *L* being the length of the latent vector (10).

**Figure 2.**
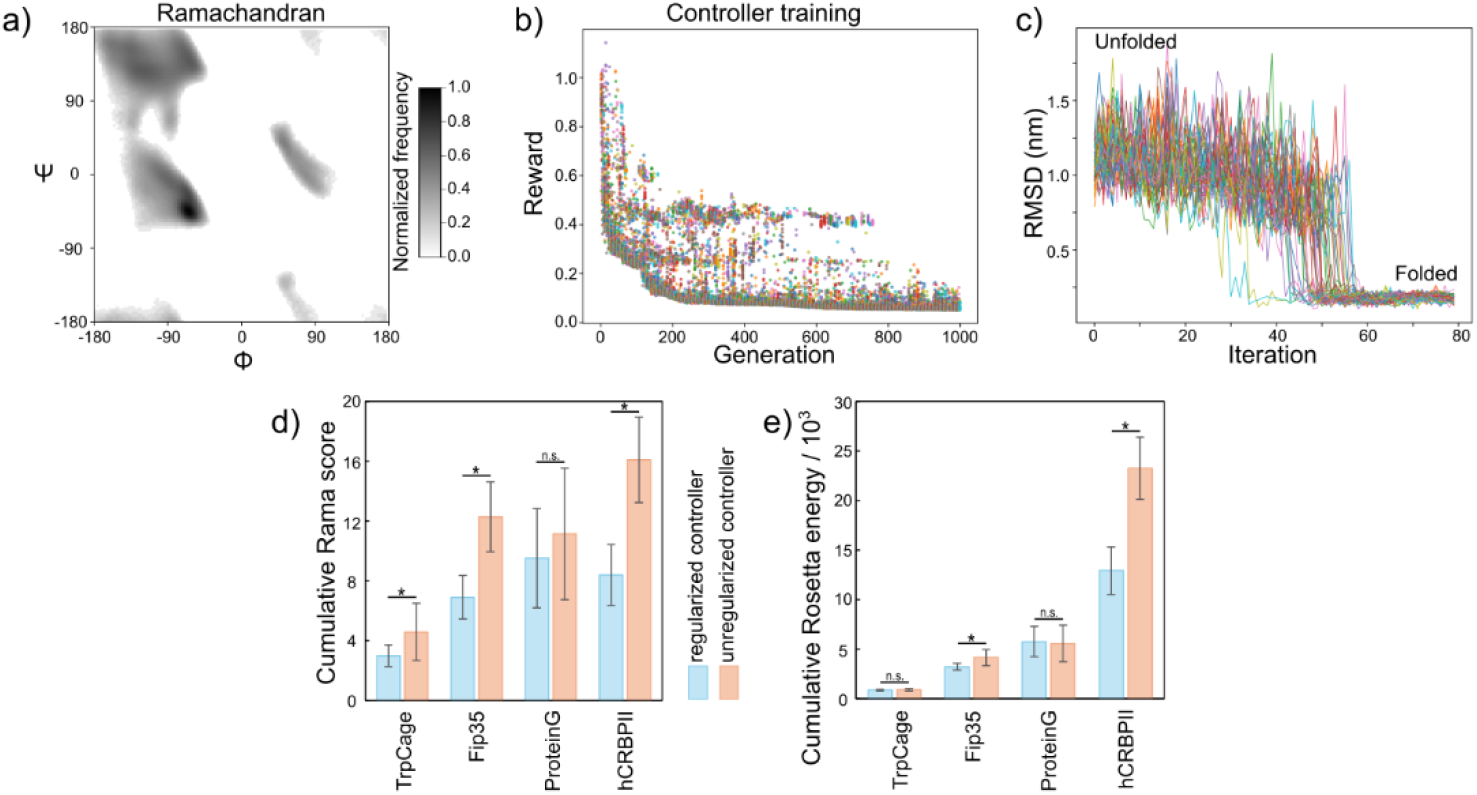
Controller training and regularization. a) Ramachandran frequency matrix *R* used to compute the regularization term. b) Training progress of the controller by CMA-ES, example of Protein G. Each generation has a population size of 20 trajectories, for a total of 1000 generations. c) RMSD of Protein G folding trajectories. Generally, we consider a conformation to be folded when its RMSD becomes < 0.3 nm. d) Cumulative Ramachandran score of the folding trajectories of the four proteins of interest. As expected, controller regularization decreases the cumulative Ramachandran score. e) Cumulative Rosetta energy calculated on the folding trajectories of the four proteins. In two cases, the controller regularization significantly reduced the cumulative Rosetta energy of the folding pathways, making them follow lower energy routes

We have used CMA-ES to evolve the controller weights for 1000 generations, with a population size of 20, starting from a different unfolded state at every generation and performing 80 or 100 moves. Controller training progress for Protein G is shown in Figure 2b. Folding trajectories for Protein G are shown in Figure 2c.

As expected, the regularization term reduced the cumulative Ramachandran score of the folding trajectories during inference (Figure 2d) (except for the case of Protein G which was not significant). Interestingly, in two cases (Fip35 and hCRBPII), it also significantly reduced the cumulative Rosetta energy score of the trajectories, indicating that the folding pathways are following more energetically favored routes (Figure 2e).

### Computation of folding pathways

Folding trajectories are computed by starting from randomly generated unfolded states and using the trained controller to iteratively perform 80 or 100 steps. We have computed 50 trajectories for each protein. Figure 2c shows the RMSD of trajectories computed for Protein G. Figure 3 shows the folding mechanism and the energy landscape (Rosetta energy score) for our proteins of interest. The folding proceeds from regions of high energy (unfolded) towards regions of low energy, with the native state always being the basin with the lowest energy. The energy of the protein configurations has been obtained using the default full atom score function of Rosetta (ref2015), which is a weighted sum of energy terms that represent physical forces and statistical terms, measured in Rosetta Energy Units (REU^22^). While it does not correspond to the thermodynamic energy landscape of the protein (because it lacks entropic contributions), it still contains valuable information about the relative stability between the conformations, the folding pathways, and the structural intermediates.

**Figure 3.**
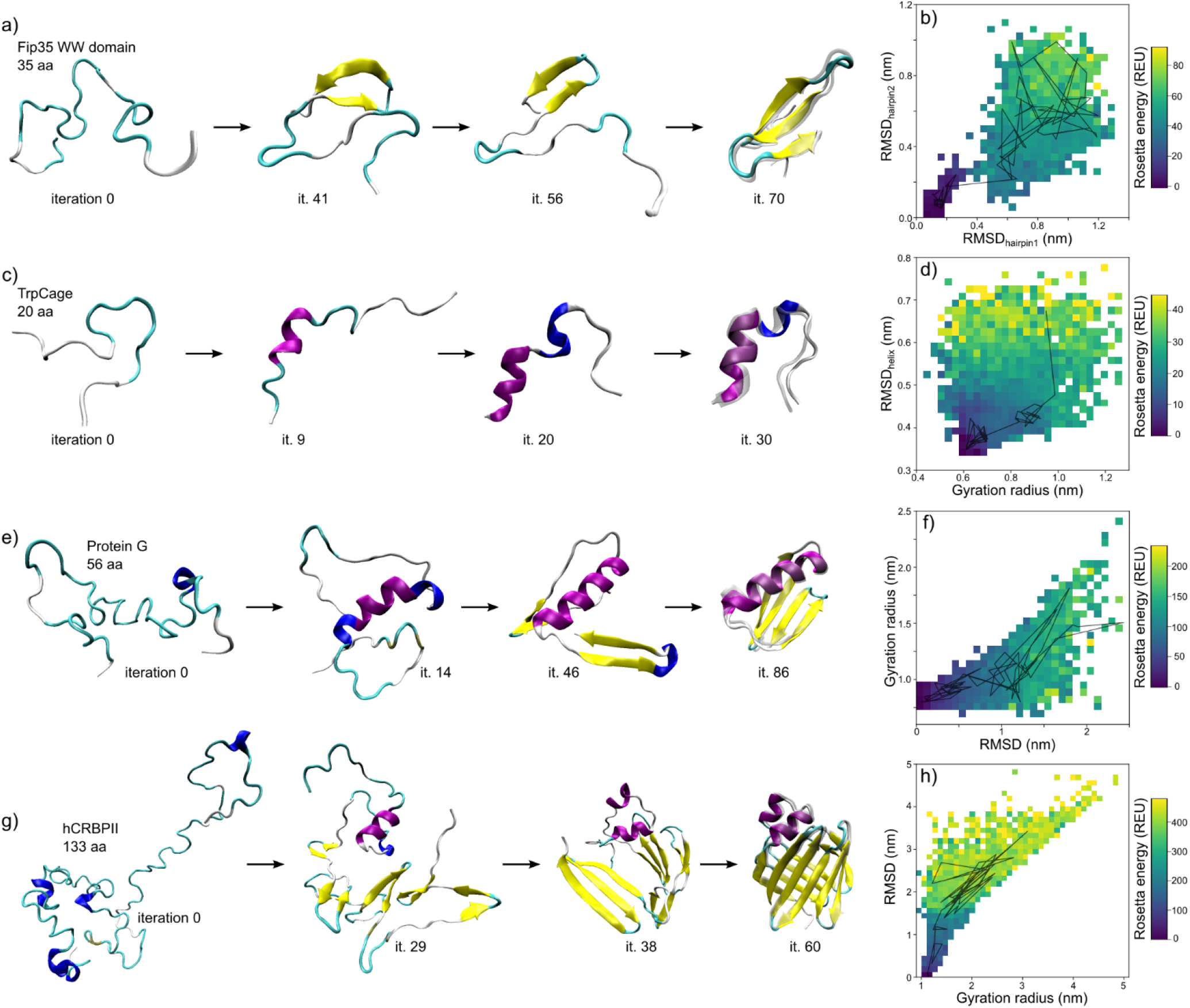
Inference of protein folding pathways. Folding mechanism and Rosetta energy landscape as a function of various collective variables for Fip 35 WW domain (a-b), TrpCage (c-d), Protein G (e-f), hCRBPII (g-h). One single folding trajectory of choice is drawn on top of each energy landscape. The energy is given in Rosetta Energy Units (REU), which are not directly transferable into a physical unit of measure but can be used to compare the relative energy of different conformations of the same protein.

### Validation

To measure the accuracy of the folding pathways generated by the World Model, we computed their path similarity^28,29^ to available data from long equilibrium MD (DE Shaw Research^30^) and other trajectories from different enhanced path sampling methods: the so-called Bias Functional scheme^10,31^ (an enhanced path sampling method that generates folding pathways using ratchet-and-pawl MD and then selects the least biased trajectories), and the “Discard-and-Restart” MD method^9^ (an efficient algorithm to compute folding pathways by repeatedly running short simulations and discarding those that do not fold forward). Results are shown in Figure 4. Further details about the calculation of the path similarity can be found in the supplementary section 3.

**Figure 4.**
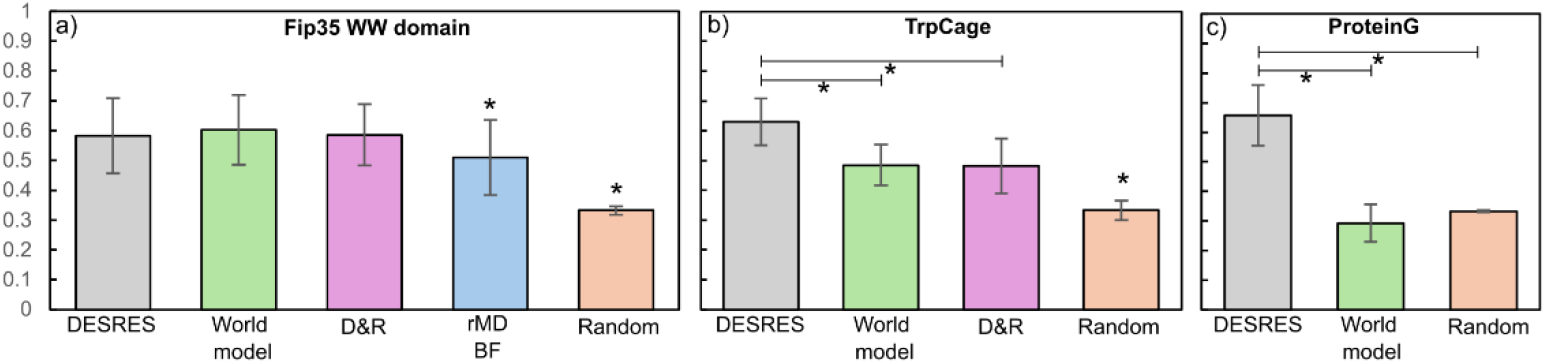
Path similarity to equilibrium MD. Path similarity analysis for Fip35 WW domain (a), TrpCage (b) and Protein G (c). DESRES: David E Shaw Research; D&R: Discard and Restart algorithm; rMDBF: Ratcheted MD Bias Functional. Asterisk indicates that the difference is statistically significant. When the asterisk is alone, it indicates statistical significance against all the other comparisons in the same plot. The same analysis was not possible for the protein hCRBPII due to unavailability of equilibrium MD data.

Our trajectories of the Fip35 WW domain have a high path similarity to equilibrium MD trajectories (Figure 4a). The order of formation of its native contacts is indistinguishable from the equilibrium trajectories and more accurate than the rMDBF method. For what concerns the TrpCage miniprotein, accuracy is a bit less and does not outperform the D&R method (Figure 4b). Moreover, the order of formation of native contacts for TrpCage does not entirely follow the folding mechanism at equilibrium. Finally, accuracy is very low for Protein G, where the path similarity indicates an order of formation of native contacts close to randomness and very different from equilibrium simulations (Figure 4c). These data indicate that this generative model is not always successful at predicting accurate folding pathways. By learning random actions from the rollout data and by minimizing the energetic cost of the trajectories, the model seems to be somewhat effective in capturing the folding dynamics of small proteins (e.g. Fip35 and TrpCage) which have a more restricted conformational space. For bigger proteins (e.g. ProteinG), this strategy is probably not enough, and the moves generated by the controller still resemble the randomness learned from the rollout data and, by consequence, the folding pathways follow a random order of native contact formation.

The method is able to generate pathways connecting two structural states (e.g. unfolded to native). The path similarity results indicate that such conformations might sometimes be artificial and not follow the equilibrium distribution. However, it is possible to reconstruct (up to a certain degree) the underlying unbiased dynamics of the system by performing local relaxations of the structures along a predefined collective variable (e.g. RMSD to native), by means of running short MD simulations^31^. In contrast to extracting slow properties from long MD simulations, Markov State Models (MSM) can construct the dynamics from short simulations starting from different conformations^32,33^. In MSM, simulations only need to reach local equilibrium (which is achieved in a few nanoseconds^31^) within individual substates and eventually transition to neighboring ones. The short unbiased trajectories give the conditional transition probabilities to start from the RMSD bin *r* and end up in bin *r′*. The goal is to measure the probability that, after some short time *Δt*, you find yourself in bin *r’*. This allows you to build the *transition matrix T*:

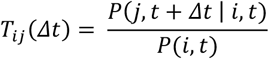

where *P*(*j*, t + *Δ*t | *i*, t) is the probability of being in state *j* at time *t*+*Δt* given that the system was in state *i* at time *t*, and *P*(*i*, t) is the probability of the system being in state *i* at time *t*. Each row of the transition matrix is then normalized to represent a conditional probability distribution ∑_*j*_ *T*_*ij*_ (*Δ*t) = 1. The stationary probability distribution then corresponds to the eigenvector corresponding to eigenvalue 1. *Δt* has been chosen to be 2ns, which is on the same timescale as the local thermalization of metastable states i.e. the degrees of freedom orthogonal to a collective variable (such as Q, the fraction of native contacts, or RMSD to native)^31^. Moreover, it also corresponds to the frequency of which coordinates have been printed in the reference trajectories from DESRES.

For each protein, we have divided the RMSD space in 50 bins and extracted a structure representing each bin. Then, we have run 40ns of MD in explicit solvent (protocol in the Material and Methods). A cartoon scheme of the procedure is shown in Figure 5a. This reconstruction method works under the assumption that even if the original trajectories were biased, these short runs (done without any bias) reflect how the system truly evolves locally. It also assumes that we can treat the dynamics as a Markov chain on the RMSD bins (i.e., the future state depends only on the current bin not on the full history; and the microstates represent separated regions of the state space).

**Figure 5.**
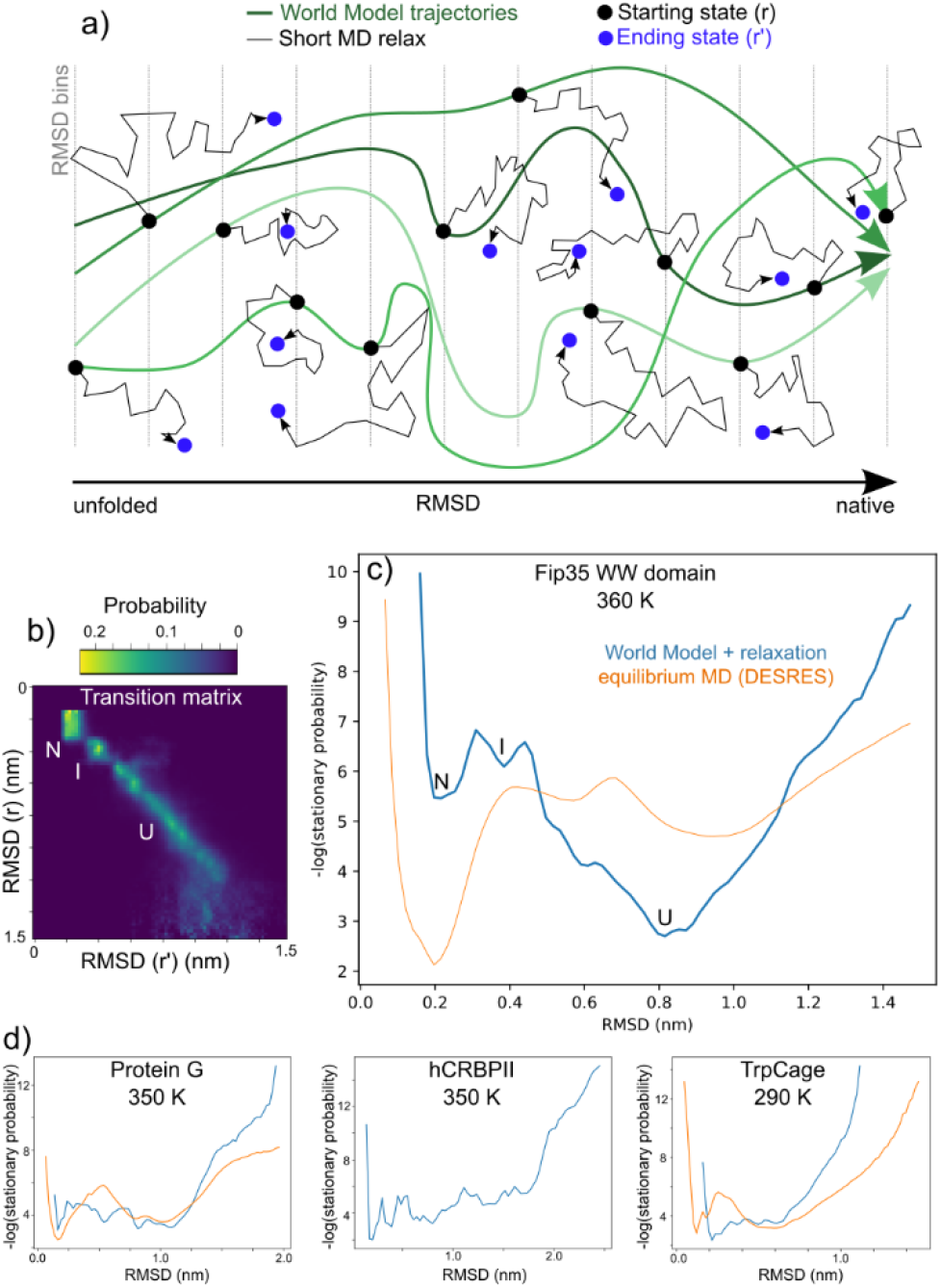
Reconstruction of unbiased dynamics by Markov State Modeling. a) Cartoon scheme of the MSM approach for the reconstruction of the true dynamics. Short MD relax simulations are started from RMSD bins from randomly chosen World Model trajectories. b) Transition matrix of the MD relaxation for the Fip35 WW domain at 360K (total run time 40ns, Δt = 2ns). c)Reconstruction of the folding landscape for Fip35 WW domain. World model after relaxation is shown in blue, equilibrium MD from DESRES is shown in orange. The landscape identified a 3-state folding mechanism: native (N), intermediate (I) and unfolded (U). d)Folding landscape for Protein G, hCRBPII and TrpCage, respectively. No equilibrium data are available for hCRBPII.

Figure 5b shows the transition matrix for Fip35 WW domain, and the reconstruction of the stationary probability is shown in 5c. The shape of the distribution is not identical to equilibrium MD, but both methods identified 3 basins corresponding to a 3-state folding (native, intermediate and unfolded). With some discrepancies, the method was able to generally reconstruct the folding landscape of our proteins, with a shape and magnitude reasonably similar to equilibrium MD (Figure 5c-d). The discrepancies might arise from different MD protocols and from the fact that the process of local relaxation cannot be entirely described as a Markovian process by a single collective variable such as RMSD, as it might fail to capture the conformational separation of states.

## CONCLUSIONS

We have demonstrated that generative World Models for policy evolution in the latent space can be implemented in the context of biomolecular simulations. Folding policy evolution becomes possible in the latent space, overcoming the need of high computational resources necessary to compute the high-dimensional real environment. We successfully computed the folding pathway for proteins of variable sizes, including a medium-large protein (133 aa) whose folding dynamics would be inaccessible by standard simulation methods. We show that structure-based regularization terms (Ramachandran score) can be used to influence the folding dynamics by reducing the energetic cost of the trajectories.

The method still needs to be improved, as it does not always predict accurate folding pathways. It can be coupled with equilibrium MD to restore the real dynamics of the system, up to a certain degree, under the assumptions of MSM. However, this is the first application of generative World Models in the context of biomolecular simulations, and we believe it can be further improved to enhance its structural accuracy. Future developments of the method could also include its generalization to perform RNA folding simulations on their dihedral angles (α, β, γ, δ, ε, ζ)^34^ or their simplifications (pseudotorsions η, θ)^35^. Additionally, we plan to design the World Model to perform unguided biomolecular simulations (i.e. without the need to specify the target), by encoding the biomolecule’s energy into the latent space, in order to be able to predict the energy of future configurations. The controller can then be trained to yield lower energy structures which ultimately lead to the native in an unguided manner.

## METHODS

The World Model has been built via custom-made code written in Python, using Keras and tensorflow to build the neural networks. Training data and structural calculations on the proteins have been made using pyRosetta, the Python-based interface to the Rosetta molecular modeling tool. Controller training (CMA) has been performed using the Python cma library.

Molecular dynamics simulations have been performed using GROMACS. Protein structures were positioned in a dodecahedral box with 12 Å minimum distance from the box walls. Water molecules have been modeled with TIP3P water and charges neutralized with Na^+^ and Cl^-^ counterions. Each system was then energy minimized by steepest gradient descent (5000 steps), followed by NVT equilibration (500 ps) and NPT equilibration (500 ps). Temperature was chosen to match available MD data and is shown in Figure 5. MD was run at the same temperature as the equilibrations and was performed using the leap-frog integrator with a dt of 2 fs for 20’000’000 steps (total 40ns). Non-bonded interactions such as Van-der-Waals and Coulomb were set with a cutoff of 1.6 nm and have been computed on CPU. Particle Mesh Ewald was employed for long range electrostatics and was computed on GPU. Parrinello-Rahman barostat was used for pressure coupling of the system, and Nose-Hoover thermostat for temperature coupling. GROMACS protocol files (.mdp files) for energy minimization, for NVT/NPT equilibration and for MD can be found in the github repository.

MD calculations have been run on the High Performance Computing facility of Yale University. The World Model was built and trained on a local Mac Pro from 2013.

## ASSOCIATED CONTENT

### Supporting information

More information are available in the supplementary material.

## AUTHOR INFORMATION

### Author contributions

A.I. developed the models, designed the study, wrote the code, analyzed the data. E.C. and J.I. provided programming and data analysis support. M.G supervised the study and determined the scientific workflow. M.G. and A.I. wrote the manuscript.

### Notes

The authors declare no competing financial interests.

