## supplementary material for "Generative World Models to compute protein folding pathways"

**Content:**

1. **Models' size and architecture**
2. **Calculation of the path similarity between trajectories**
3. **Decoding and reconstruction of protein configurations**
4. **The actions on the dihedrals**
   **References**

### Abbreviations

VAE, variational autoencoder; FFN, feed-forward network; CMA-ES, covariance matrix adaptation evolution strategy; MD, molecular dynamics; RMSD, root mean squared displacement; D&R, discard and restart; AI, artificial intelligence.

### 1. Models' size and architecture

Variational Autoencoder 1 (VAE1): The first variational autoencoder takes Φ, Ψ and the $C_\alpha$ distances as input and encodes them into a latent vector $z_{features}$ of reduced dimensionality (length 10). The number of parameters of the model therefore depends on the number of residues of the protein.

Variational Autoencoder 2 (VAE2): The second variational autoencoder takes the action on Φ (ΔΦ) and Ψ (ΔΨ) and encodes them into a latent vector $z_{action}$ of reduced dimensionality (length 10). Also in this case, the number of parameters of the model depends on the number of residues of the protein. The architecture of VAE1 and VAE2 is the same.

Feed Forward Network (FFN): Predicts the next latent vector ($z_{features}(t+1)$) given the current latent configuration ($z_{features}(t)$) and latent action ($z_{action}(t)$). It is a series of expanding and contracting Dense layers. The number of parameters of the model does not depend on the protein and is therefore the same for every protein.

Controller: Predicts the latent action ($z_{action}$) to be performed on the latent configuration ($z_{features}$) in order to drive it towards the target state.

| Model | # params |
|---|---|
| VAE1 TrpCage | 11'960 |
| VAE2 TrpCage | 5'580 |
| VAE1 Fip35 | 14'685 |
| VAE2 Fip35 | 6'896 |

| Model | # params |
|---|---|
| VAE1 Protein G | 36'980 |
| VAE2 Protein G | 16'820 |
| VAE1 hCRBP II | 203'619 |
| VAE2 hCRBP II | 90'976 |

| Model | # params |
|---|---|
| FFN | 7'770 |
| Controller | 220 |

**Table S1.1: Size of the models.**

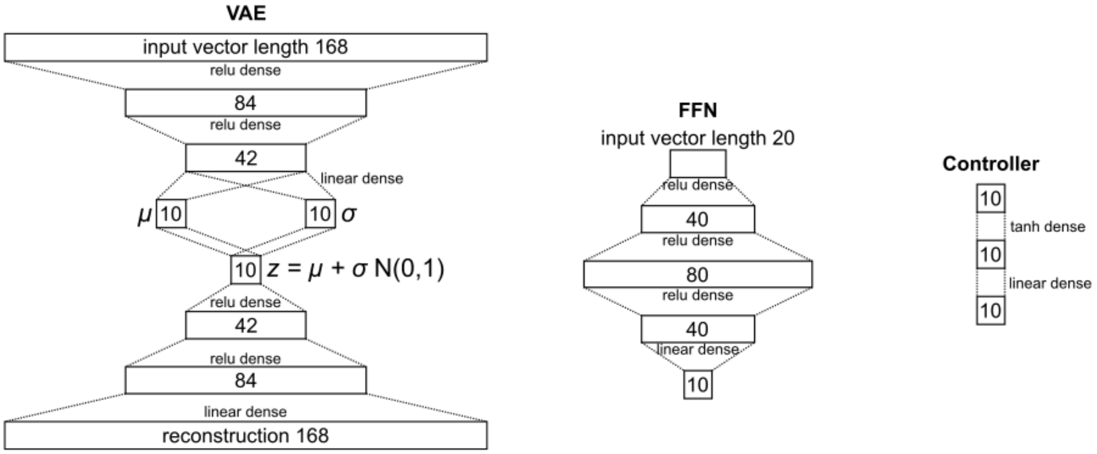


**Figure S1.1: Architecture of the models.** The VAE shown here corresponds to VAE1 Protein G.

### 2. Calculation of the path similarity between trajectories

Path similarity has been computed as indicated in [1,2]. It quantifies how consistently native contacts form in the same temporal order across two given pathways. This parameter ranges from 0, indicating no similarity, to 1, indicating that all native contacts form in exactly the same sequence for both trajectories. To calculate this quantity, we define a matrix *M* that represents the order of native contact formation between atoms (Cα). Denoting the formation time of the i-th native contact in the k-th trajectory, the matrix element for the k-th pathway is given by:

$$M_{ij}(k) = \begin{cases} 1 \text{ if } t_{ik} < t_{jk} \\ 0 \text{ if } t_{ik} > t_{jk} \\ \frac{1}{2} \text{ if } t_{ik} = t_{jk} \end{cases}$$

The path similarity *s* between trajectories *k* and *k'* is then defined as:

$$s(k, k') = \frac{1}{N_C(N_C - 1)} \sum_{i \neq j} \delta(\, M_{ij}(k) - M_{ij}(k')\,)$$

where $\delta$ is the Dirac delta function and *Nc* is the total number of native contacts.

### 3. Decoding and reconstruction of protein configurations

From the latent configurations $z_{features}$, the decoder of VAE1 could be used to obtain back the values of Φ, Ψ and the $C_\alpha$ distances from native. A specific combination of Φ, Ψ dihedral angles corresponds to a unique protein backbone conformation[3]. However, the slight noise derived after decoding (intrinsic of Variational Autoencoders[4]) and the model's reconstruction error causes dihedral differences to cumulatively propagate across the protein chain and rotate the backbone in an undesired manner. The value of the $C_\alpha$ distances from native was used to preserve the correct backbone orientation and correct for the noise and reconstruction error of the VAE1. The objective is therefore to minimize the discrepancy between the calculated and reference $C_\alpha$ distances by slightly adjusting dihedral angles (Φ and Ψ). This was achieved by minimizing the following objective function *E*:

$$E(\Phi, \Psi) = \sqrt{\frac{1}{N} \sum_{i=1}^{N} \left( d_{calc}(i;\, \Phi, \Psi) - d_{ref}(i) \right)^2}$$

where: $d_{calc}(i;\, \Phi, \Psi)$ corresponds to the $C_\alpha$ distance deviation for the current configuration, computed using the dihedral angles (Φ and Ψ); $d_{ref}(i)$ corresponds to the reference $C_\alpha$ distance deviation (from the decoded vector), *N* is the total number of residues of the protein. The optimization attempts to find dihedral angles Φ and Ψ such that $\min_{\Phi,\Psi} E(\Phi, \Psi)$. The minimization was performed using the BFGS algorithm[5], updating the angles to bring the calculated $C_\alpha$ distances closer to the reference. The optimization was stopped either when the RMSD between the calculated and reference Cα distances went below a threshold (0.35) or the iteration count exceeded a set limit (75000). After the optimization process, an energy minimization step was performed using pyRosetta[6].

### 4. The actions on the dihedrals

During the creation of the rollout data, a new configuration is generated by performing a random rotation of the dihedrals on the previous configuration. First of all, we define the magnitude (*m*) of the action by picking a random number from a uniform distribution between 0° and 30°. This step defines how strong the action on the dihedrals is gonna be overall for that step. Then, for each dihedral angle Φ and Ψ, we pick a random number from a uniform distribution between -*m* and +*m*. This defines the rotation to be performed on that particular dihedral angle.

It is therefore clear that the dihedral actions at each step are of complete random nature, and barely preserve an underlying pattern that is learnable by an AI model. This is indeed the case for our VAE2, which maintains a high reconstruction error during training and inference, as shown in Figure S3 (the VAE2 is able to learn the magnitude *m* of the action, but not always the value of the individual dihedral moves).

However, this does not defy the purpose of VAE2. Mapping the actions on a continuous latent space (a multivariate Gaussian distribution) guarantees a smooth interpolation between the points[7]. This means that during its training, the FFN builds its own understanding of how the latent action ($z_{action}$) affects the dihedrals moves ($z_{features}$), and predicts the next structure ($z_{features}(t+1)$). Then, during policy evolution, the controller is going to evolve its weights in order to predict the next action ($z_{action}$) based on the FFN's understanding of the world (World Model) i.e. the structural transitions. However, it is important to note that the action loses its meaning in the real environment and, after decoding, does not relate to a realistic action anymore. The real action performed on the structure at any particular step can anyway be inferred later by measuring the dihedral differences before and after the move.

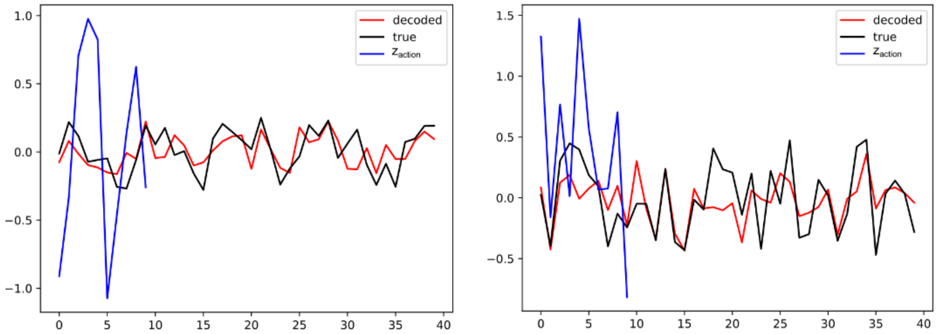


**Figure S4.1: Examples of action encoding and reconstruction.** The reconstruction of the action (red line) maintains a high reconstruction error due to the stochastic nature of the action in the training data (true action in black). The blue line shows the latent action ($z_{action}$). This is an example made with the TrpCage validation data. 0 to 19 corresponds to ΔΦ, 20 to 39 corresponds to ΔΨ. The action is normalized between -1 and +1.
